# *Wolbachia* both aids and hampers the performance of spider mites on different host plants

**DOI:** 10.1101/344143

**Authors:** Flore Zélé, Joaquim L. Santos, Diogo Prino Godinho, Sara Magalhães

## Abstract

In the last decades, many studies had revealed the potential role of arthropod bacterial endosymbionts in shaping the host range of generalist herbivores and their performance on different host plants, which, in turn, might affect endosymbiont distribution in herbivores populations. We tested this by measuring the prevalence of endosymbionts in natural populations of the generalist spider mite *Tetranychus urticae* on different host plants. Focusing on *Wolbachia*, we then analysed how symbionts affected mite life-history traits on the same host-plants in the laboratory. Overall, the prevalences of *Cardinium* and *Rickettsia* were low, whereas that of *Wolbachia* was high, with the highest values on bean and eggplant and the lowest on purple, tomato and zuchini. Although most mite life-history traits were affected by the plant species only, *Wolbachia* infection was detrimental for egg hatching rate on purple and zucchini, and led to a more female-biased sex ratio on purple and eggplant. These results suggest that endosymbionts may affect the host range of polyphagous herbivores, both by aiding and hampering their performance, depending on the host plant and on the life-history trait that affects performance the most. Conversely, endosymbiont spread may be facilitated or hindered by the plants on which infected herbivores occur.

## INTRODUCTION

Although generalist herbivores are able to colonize several host plants, their performance on different host plants is variable. Whereas some studies suggest that the host range of herbivores is mostly determined by geographical location (Calatayud *et al.*, 2016), others suggest that this range is determined by host-plant nutritional quality (Schoonhoven *et al.*, 2005) or host-plant defences (Becerra, 1997). Still, the proximate mechanisms allowing populations to colonize particular host plants remain elusive.

Herbivores harbour a rich community of microorganisms, ranging from their gut microbiota and intracellular vertically-transmitted endosymbionts to plant bacteria and viruses of which they serve as vectors, and there is growing evidence of the impact of such communities on herbivore performance on plants (Hosokawa *et al.*, 2007, Clark *et al.*, 2010, Frago *et al.*, 2012, Hansen & Moran, 2014, Oliver & Martinez, 2014, Zhu *et al.*, 2014, Shikano *et al.*, 2017). Obvious candidates to influence plant colonization by herbivorous arthropods are their heritable endosymbionts (Clark *et al.*, 2010, Feldhaar, 2011, Ferrari & Vavre, 2011, Frago *et al.*, 2012, Jaenike, 2015). Due to their vertical mode of transmission, the fitness of such symbionts is tightly linked to that of their host and they are likely to benefit their host in order to increase their own transmission (Fine, 1975). Indeed, endosymbionts have been shown to affect the host-plant range of herbivorous arthropods (Hosokawa *et al.*, 2007, Tsuchida *et al.*, 2011, Sugio *et al.*, 2015, Wagner *et al.*, 2015, Giron *et al.*, 2017) or to increase performance on certain plant species (Wilkinson *et al.*, 2001, Leonardo & Muiru, 2003, Ferrari *et al.*, 2004, Tsuchida *et al.*, 2004, Ferrari *et al.*, 2007, Hosokawa *et al.*, 2007, Su *et al.*, 2013, Su *et al.*, 2015, Wagner *et al.*, 2015), while decreasing performance on others (Chen *et al.*, 2000, Leonardo & Muiru, 2003, Ferrari *et al.*, 2007, Chandler *et al.*, 2008, McLean *et al.*, 2011, Wagner *et al.*, 2015). In some cases, increased host performance is due to endosymbionts acting as nutritional mutualists, directly supplying their arthropod hosts with nutrients or enzymes that are missing in their plant diet (reviewed by Chaves *et al.*, 2009, Douglas, 2009), or displaying compensatory effects during periods of nutritional deficiency (Su *et al.*, 2014). Finally, endosymbionts may also enable arthropods to manipulate phytohormonal profiles (Kaiser *et al.*, 2010, Body *et al.*, 2013), resource allocation (Hackett *et al.*, 2013), and anti-herbivory defences (Barr *et al.*, 2010, Su *et al.*, 2015). Conversely, symbiont-mediated decreased host performance on particular plants might be due to the nutrient profile (e.g., specific amino acids and nitrogen content) of these plants, which promotes deleterious symbiont traits and disturbs the host control over bacterial abundance (Wilkinson *et al.*, 2007, Chandler *et al.*, 2008).

Such variable effects of endosymbionts on herbivore plant use may contribute to variation in the abundance and distribution of herbivorous arthropods (Douglas, 2009, Hansen & Moran, 2014). Conversely, as symbiont-herbivore interactions may differ according to the host plant, and nutrition of herbivore host can affect the within-host symbiont density (Wilkinson *et al.*, 2001, Wilkinson *et al.*, 2007, Chandler *et al.*, 2008, Zhang *et al.*, 2016), the host plant can also affect endosymbiont distribution in the field (Leonardo & Muiru, 2003, Simon *et al.*, 2003, Ferrari *et al.*, 2004, Tsuchida *et al.*, 2004, Chandler *et al.*, 2008, Ahmed *et al.*, 2010, Brady & White, 2013, Pan *et al.*, 2013, Guidolin & Consoli, 2017). However, most studies addressing these questions have been conducted on sap-feeding insects and whether symbiont prevalence and their effects on their herbivorous host vary with the host plant remains unstudied in other systems.

The two-spotted spider mite *Tetranychus urticae*, a cosmopolitan agricultural and horticultural pest that feeds on cell content, is a highly polyphagous arthropod, feeding on more than 1100 plant species (Migeon & Dorkeld, 2006-2017). This generalist herbivore rapidly adapts to novel host plants (Fry, 1990, Agrawal, 2000, Magalhães *et al.*, 2007), sometimes forming host races (Magalhães *et al.*, 2007), and may harbour several endosymbiontic bacteria with variable prevalence among populations (Enigl & Schausberger, 2007, Gotoh *et al.*, 2007, Staudacher *et al.*, 2017). Among them, *Wolbachia* is the most prevalent (Liu *et al.*, 2006, Gotoh *et al.*, 2007, Ros & Breeuwer, 2009, Zhang *et al.*, 2016, Zélé *et al.*, 2018) and induces variable fitness effects in spider mites. For instance, it can decrease (Perrot-Minnot *et al.*, 2002, Suh *et al.*, 2015), not affect (Breeuwer, 1997, Vala *et al.*, 2000, Perrot-Minnot *et al.*, 2002, Vala *et al.*, 2002, Gotoh *et al.*, 2007), or increase (Vala *et al.*, 2002, Gotoh *et al.*, 2007, Xie *et al.*, 2011) their fecundity. Given these variable effects, it is as yet unclear whether *Wolbachia* will facilitate or hamper host-plant colonization by spider mites.

Here, we measured the prevalence of the three most prevalent endosymbionts of *T. urticae*, namely *Wolbachia*, *Cardinium*, and *Rickettsia*, on five different host plants in Portugal. Subsequently, we explored whether the effect of *Wolbachia* on the performance of *T. urticae* hinges on the plant that is being colonized. Finally, we discuss the importance of possible mechanisms leading to our results as well as the potential adaptive significance of the presence of *Wolbachia* for plant colonization by *T. urticae*.

## MATERIALS AND METHODS

### Effect of the host plant on endosymbiont prevalence in the field

To determine whether the prevalence of *Wolbachia, Cardinium* and *Rickettsia* in natural *T. urticae* populations varied with the host plant, spider mites were collected on bean (*Phaseolus vulgaris*, Fabaceae), eggplant (*Solanum melongena*, Solenaceae), purple morning glory (*Ipomoea purpurea*, Convolvulaceae, hereafter “purple”), zucchini (*Cucurbita pepo*, Cucurbitaceae), and tomato (*Solanum lycopersicum*, Solenaceae) across 12 different locations (Table 1). These plants were selected because they are part of the natural host range of *T. urticae* but belong to different families. Sampling sites consisted of open fields, greenhouses or organic vegetable gardens, while being insecticide/pesticide free to avoid this potential confounding effect. Infested leaves were detached and placed in closed plastic boxes that were brought to the laboratory. On the same day, 50 adult females were haphazardly picked from each population and their species determined at the individual level based on morphological characteristics under a binocular microscope. These females were then placed on 2 cm^2^ leaf discs of the same plant species on which they were found, and allowed to lay eggs for 4 days. Subsequently, 20 of these females were randomly selected and individually tested for the presence of *Wolbachia*, *Cardinium* and *Rickettsia* on entire mites without DNA extraction by multiplex PCR using genus-specific primers as described in (Zélé *et al.*, 2018). Subsequently, for each population, the DNA of a pool consisting of one daughter from each of these females was extracted, then a PCR-based method to identify the mite species was performed by multiplex PCR as described in (Zélé *et al.*, 2018). If a pool could not be assigned unambiguously to *T. urticae* (see Table S1 in Additional file 1), all data concerning endosymbiont prevalence were discarded. This process was repeated until obtaining endosymbiont prevalence data for 5 populations per plant, except for purple, for which we could obtain only 2 populations of *T. urticae* due to the weak infestation rate of this plant by this spider-mite species, and despite a large sampling effort (Table S1).

**Table 1.**
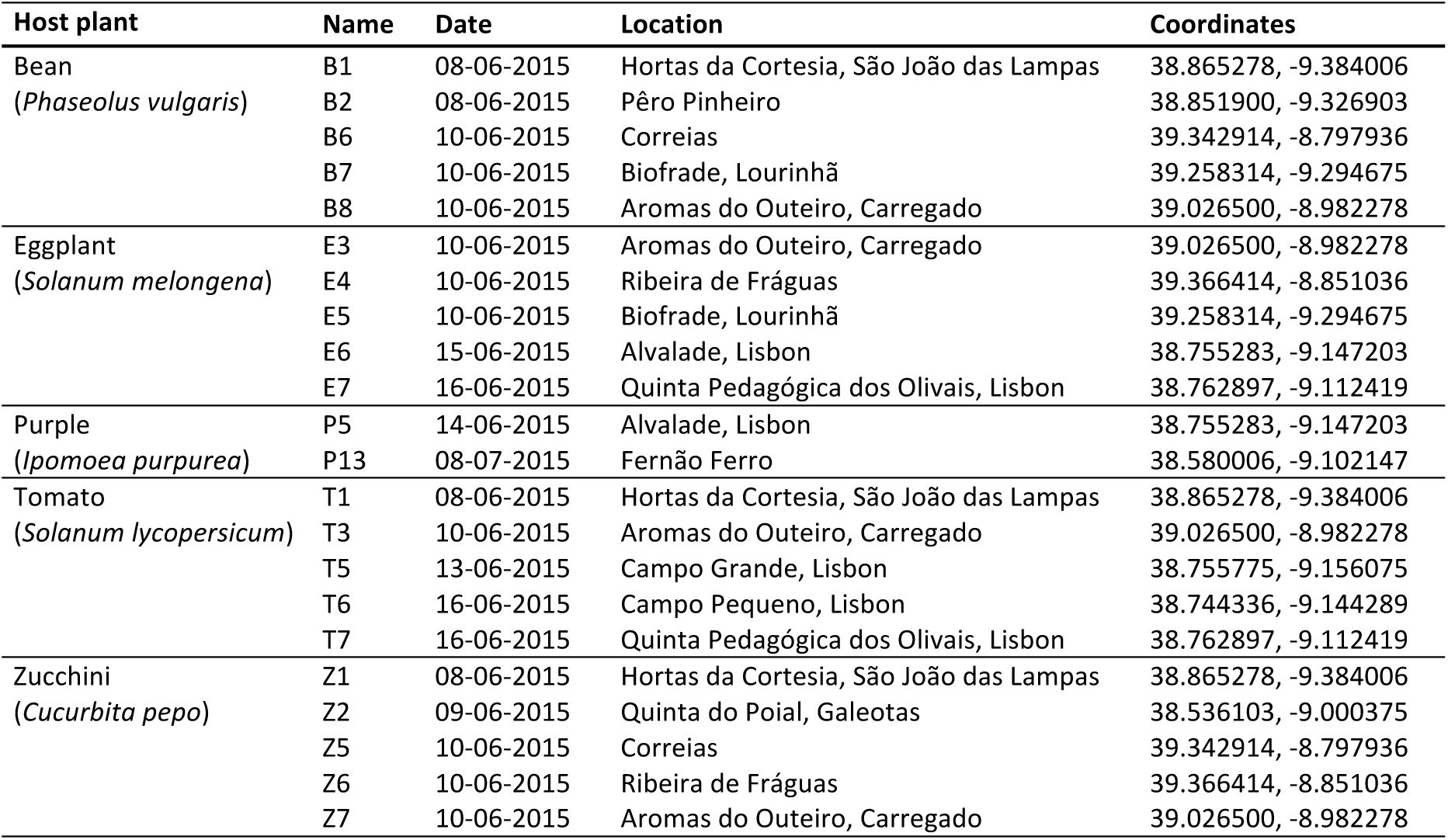
*Tetranychus urticae* populations collected on five different host plants across 12 different locations in June-July 2015 and used to study the plant effect on the prevalence of *Wolbachia*, *Cardinium* and *Rickettsia*.

### Effect of *Wolbachia*, the host plant, and their interaction on the performance of spider mites

#### Spider mite populations, tetracycline treatment and population rearing

The spider-mite population used was originally collected on *Datura* plants at Aldeia da Mata Pequena, Portugal, in November 2013 and kept in a mass-rearing environment (>5 000 individuals) on bean plants (var. *Enana*), under controlled conditions (25°C, photoperiod of 16L:8D) since then. This population, hereafter called Wi, was found uninfected by *Rickettsia*, *Spiroplasma* or *Arsenophonus* but fully infected by *Wolbachia* in the field (Zélé *et al.*, 2018). Although this population was also slightly infected by *Cardinium* (Zélé *et al.*, 2018), this endosymbiont has been rapidly lost following laboratory rearing (unpublished data). To obtain a *Wolbachia*-uninfected (Wu) population with a similar genetic background, roughly 3 months after collection 30 adult females of the Wi population were placed in petri dishes containing bean leaf fragments placed on cotton with a tetracycline solution (0.1 %, w/v). This treatment was applied continuously for three successive generations (Breeuwer, 1997), then the population was maintained in a mass-rearing environment without antibiotics for c.a. 12 generations before the experiment to avoid (or limit) potential side effects of the antibiotic treatment (e.g. O’Shea & Singh, 2015) and allow mites to recover potential loss of gut. Before use, up to 20 individual females and pools of 100 females were checked by PCR to confirm the absence and presence of *Wolbachia* infection in Wu and Wi populations, respectively.

#### Performance of Wolbachia-infected and uninfected females on different host plant

To determine the effect of *Wolbachia* infection and of the host plant, as well as their possible interaction, on the performance of *T. urticae*, we measured life history traits of individuals from Wi or Wu populations when placed on the same plant species as those from which mites were collected in the field study (bean: var. *Enana*, eggplant: var. *Larga Morada*, purple: var. *Vigorous*, zucchini: var. *Bellezza Negra*, and tomato: var. Money Maker). To control for age, 100 females were allowed to lay eggs for three days on detached bean leaves placed on water-soaked cotton, and the adult females resulting from those eggs were used in the experiments. Fifty mated females (10-13 days old) were haphazardly picked from either Wi or Wu cohorts and placed individually on a 2 cm^2^ leaf disc from one of the 5 different host plants. The replicates were distributed along 5 temporal blocks (10 replicates per treatment per day during 5 consecutive days). Females that were alive after 3 days were transferred to new leaf discs where they could lay eggs for another 3 days. Their survival (S) and the proportion of drowned females in the water-soaked cotton (i.e. accidental death of females trying to escape the leaf discs; PD) were followed daily during six days. The fecundity of each female was measured at days 3 and 6 and the average female daily fecundity was estimated taking into account their daily mortality (DF = total number of eggs laid per female / number of days the female was alive). The number of unhatched eggs was counted 5 days later (i.e. days 8 and 11, respectively) to estimate the hatching rate (HR = hatched eggs / total number of eggs). Adult offspring (F1 females + F1 males) was counted after 6 additional days (i.e. days 14 and 17, respectively) and used to estimate juvenile mortality (JM = [total number of eggs - number of unhatched eggs - number of F1 adults]/ total number of eggs), F1 sex ratio (SR = number of F1 males/number of F1 adults) and the number of viable offspring (VO = total number of adult offspring per female per treatment observed at the end of the experiment on each plant). The entire experiment was repeated three months later (hereafter called blocks 1 and 2) except for replicates involving tomato plants. Indeed, given a very high proportion of drowned females (88 ± 3.3 %; data not shown) and because the surviving females laid on average less than 1 egg per day (0.32 ± 0.05; data not shown) on this plant, subsequent traits could not be measured and we decided to exclude it from this experiment.

### Statistical analyses

Analyses were carried out using the R statistical package (v. 3.3.2). The different statistical models built to analyse the effect of host-plant on endosymbiont prevalence in field-collected spider-mite populations and the effects of *Wolbachia* on different host plants are described in the electronic supplementary material (Additional file 1), Table S2.

To analyse the effect of host plants on endosymbiont prevalence in field-collected mites, the prevalence of *Wolbachia* (model 1), *Cardinium* (model 2) and *Rickettsia* (model 3) were fit as binary response variables, the host plant on which mites were collected as fixed explanatory variable, and the location as random explanatory variable. Because of quasi-complete separation of some of our data, which usually causes problems with estimated regression coefficients, analyses were conducted using a mixed model bglmer procedure (blme package) with a binomial error distribution (Pasch *et al.*, 2013). When the variable “plant” was significant, a stepwise *a posteriori* procedure (Crawley, 2007) to determine differences between plants was carried out by aggregating factor levels together and by testing the fit of the simplified model using a likelihood ratio test (LRT), which is approximately distributed as a χ^2^ distribution (Bolker, 2008). Because none of the mites collected in this study were singly infected by *Cardnium* or *Rickettsia*, and the prevalence of each type of coinfection was very low (cf. Results), we did not have enough statistical power to study the effect of the host plants on the prevalence of coinfections.

To analyse the effect of *Wolbachia*, the host plant, and their interaction on the performance of spider mites, the infection status of females (i.e. Wi: infected or Wu: uninfected) and the host plants tested were fit as fixed explanatory variables, whereas block and day were fit as random explanatory variables (day nested within block). Survival data (S; model 4) were analysed using a Cox proportional hazards mixed-effect model (coxme, kinship package). Hazard ratios were obtained from this model as an estimate of the difference in mortality rate (Crawley, 2007) between our control (Wi population on bean) and each of the other factor levels. PD, a binary response variable (drowned or not; model 5), was analysed using a generalized linear mixed model with a binomial distribution (glmer, lme4 package). DF, a continuous response variable (model 6) was analysed using linear mixed-effect models (lmer, nlme package). The other proportion variables HR, SR and JM (models 7, 8, and 9, respectively) were computed using the function cbind (e.g. number of hatched eggs, males, or dead juveniles *vs*. number of unhatched eggs, females, or alive juveniles, respectively). However, due to the low daily fecundity of spider mites, these variables, as well as VO (model 10) were greatly over-dispersed. One way of handling this over-dispersion is by using quasibinomial or negative binomial pseudo distributions (Crawley, 2007) but, to our knowledge, this is not possible within the usual mixed model *glmer* procedure. Thus, we used instead a mixed model glmmadmb procedure (glmmADMB package) with zero-inflated binomial error distribution for HR, SR and JM, and zero-inflated negative binomial error distribution for VO. When a statistically significant interaction between the variables “*Wolbachia*” (Wi or Wu) and “plant” was found, the effect of *Wolbachia* was analysed for each plant separately. When only the variable “plant” was significant, *a posteriori* contrasts between host plants were performed as before.

For all analyses, maximal models were simplified by sequentially eliminating non-significant terms to establish a minimal model (Crawley, 2007), and the significance of the explanatory variables was established using χ^2^-tests or *F-tests* to account for overdispersion (Bolker, 2008). The significant values given in the text are for the minimal model, while non-significant values correspond to those obtained before deletion of the variable from the model (Crawley, 2007). Full datasets are given in Additional files 2 and 3.

## RESULTS

### Effect of the host plant on endosymbiont prevalence in the field

The prevalence of *Wolbachia* was overall high (92.7 ± 1.2 %), while that of *Cardinium* (2.5 ± 0.7 %) and *Rickettsia* (2.0 ± 0.7 %) were low (Fig. 1). In addition, while 89.3 ± 1.5 % of the mites collected in this study were infected by *Wolbachia* only, none were infected by *Cardinium* or by *Rickettsia* only. 1.4 ± 0.6 % were coinfected by *Wolbachia* and *Cardinium*, 0.9 ± 0.5 % were coinfected by *Wolbachia* and *Rickettsia*, and 1.14 ± 0.5 % where infected by these three endosymbionts (see Fig. S1 in Additional file 1 for infection statuses at the individual level). The prevalence of *Wolbachia* and of *Rickettsia* were affected by the plant on which *T. urticae* females were collected (*Χ*^2^_4_=14.79, p=0.005; model 1, and *Χ*^2^_4_=12.71, p=0.01; model 3, respectively; Fig. 1). Contrast analyses revealed that the prevalence of *Wolbachia* was higher on bean and eggplant (97.0 ± 1.7 %; *contrast bean vs eggplant*: *Χ*^2^_1_=0.51, p=0.47) than on the 3 other plants (89.2 ± 2.0 %; *Contrast purple vs tomato vs zucchini*: *Χ*^2^_2_=0.39, p=0.82; *Contrast between the two groups of plants*: *Χ*^2^_1_=14.34, p=0.0002), and that of *Rickettsia* differed only on purple (12.5 ± 5.3 %) compared to all other plants (1.0 ± 0.5 %; *contrast bean vs eggplant vs tomato vs zucchini*: *Χ*^2^_3_=2.95, p=0.40; *Contrast between this group of plants and purple*: *Χ*^2^_1_=9.76, p=0.002). Finally, the prevalence of *Cardinium*, similarly to that of *Rickettsia*, tended to be higher on purple (12.5 ± 5.3 %) compared to the other plants (1.5 ± 0.6 %), but this effect was re not statistically significant (*Χ*^2^_4_=1.61, p=0.81; model 2).

**Figure 1.**
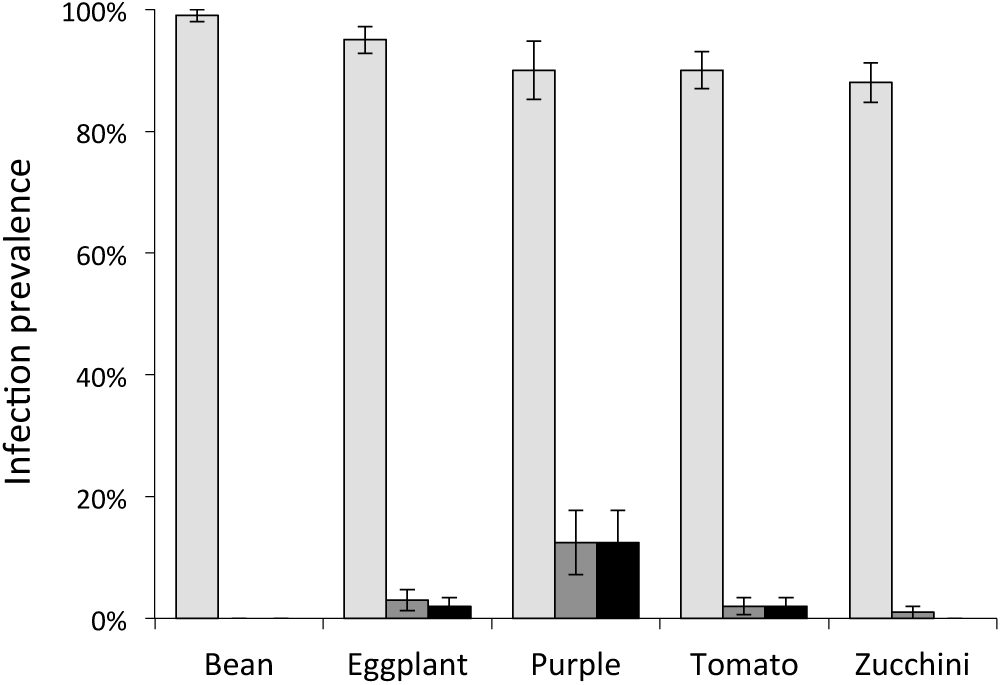
Endosymbiont prevalence in *T. urticae* females collected on different host plants. Bars represent the mean (± s.e.) infection frequencies by *Wolbachia* (light grey), *Cardinium* (dark grey), and *Rickettsia* (black) for several spider mite populations collected on bean (n=5), eggplant (n=5), purple (n=2), tomato (n=5), and zucchini (n=5).

### Effect of *Wolbachia*, the host plant, and their interaction on the performance of spider mites

Overall, there was no significant effect of *Wolbachia* (*Χ*^2^_1_= 0.73, p=0.39), of host plants (*Χ*^2^_3_= 6.84, p=0.07), or of their interaction (*Χ*^2^_3_= 3.34, p=0.34; model 4; Table 1 and Fig. S2 in Additional file 1) on survival (**S**) over the 6 first days of the experiment. However, host plants affected significantly the proportion of drowned mites (PD; *Χ*^2^_3_= 23.14, p<0.0001), regardless of *Wolbachia* infection (*Wolbachia* effect: *Χ*^2^_1_= 1.35, p=0.25; *Wolbachia*-plant interaction: *Χ*^2^_3_=0.70, p=0.87; model 5; Table 2).

**Table 2.** Effect of *Wolbachia* and of host plants on the performance of spider mites. Mean (± s.e.) values of both *Wolbachia*-infected (Wi) and uninfected (Wu) *T. urticae* on the different plants studied (bean, purple, zucchini and eggplant) are represented for each one of the performance traits measured in this study. For hatching rate, juvenile mortality and sex ratio, estimates were obtained from the GLMM statistical models and take into account variation among females, as well as the correction for zero-inflation and day within block as random effect.

| Variable of interest | Bean |  | Purple |  | Zucchini |  | Eggplant |  | Significance of explanatory variables and their interaction |  |  |
| --- | --- | --- | --- | --- | --- | --- | --- | --- | --- | --- | --- |
|  | Wi | Wu | Wi | Wu | Wi | Wu | Wi | Wu | Plant * *Wolbachia* | Plant | *Wolbachia* |
| **Log Hazard Ratio (S)** | - | 0.15 ± 0.21 | -0.03 ± 0.22 | -0.21 ± 0.31 | -0.06 ± 0.23 | -0.59 ± 0.32 | 0.18 ± 0.22 | -0.28 ± 0.30 | $\chi^2_3=3.34$, p=0.34 | $\chi^2_3=6.84$, p=0.08 | $\chi^2_1=0.88$, p=0.35 |
| **Proportion of drowned (PD)** | 0.16 ± 0.04 | 0.13 ± 0.03 | 0.26 ± 0.04 | 0.22 ± 0.04 | 0.34 ± 0.05 | 0.26 ± 0.04 | 0.34 ± 0.05 | 0.34 ± 0.05 | $\chi^2_3=0.70$, p=0.87 | $\chi^2_3=23.14$, p<0.0001 | $\chi^2_1=1.35$, p=0.25 |
| **Daily fecundity (DF)** | 4.76 ± 0.27 | 4.43 ± 0.26 | 3.54 ± 0.24 | 3.42 ± 0.21 | 3.26 ± 0.22 | 3.25 ± 0.22 | 2.32 ± 0.21 | 1.88 ± 0.15 | $\chi^2_3=1.21$, p=0.75 | $\chi^2_3=129.33$, p<0.0001 | $\chi^2_1=2.06$, p=0.15 |
| **Hatching rate (HR)** | 0.97 ± 0.01 | 0.96 ± 0.01 | 0.96 ± 0.01 | 0.98 ± 0.01 | 0.92 ± 0.01 | 0.95 ± 0.01 | 0.94 ± 0.01 | 0.93 ± 0.02 | $F_{3,697}=5.47$, p=0.001 | - | - |
| **Juvenile mortality (JM)** | 0.18 ± 0.03 | 0.20 ± 0.01 | 0.12 ± 0.01 | 0.11 ± 0.01 | 0.19 ± 0.02 | 0.16 ± 0.02 | 0.32 ± 0.02 | 0.27 ± 0.03 | $F_{3,689}=1.85$, p=0.14 | $F_{3,693}=48.23$, p<0.0001 | $F_{1,692}=0.01$, p=0.92 |
| **Sex ratio (SR)** | 0.19 ± 0.01 | 0.20 ± 0.01 | 0.21 ± 0.01 | 0.24 ± 0.02 | 0.21 ± 0.02 | 0.23 ± 0.02 | 0.17 ± 0.02 | 0.24 ± 0.03 | $F_{3,681}=2.48$, p=0.04 | - | - |
| **Viable offspring (VO)** | 19.03 ± 1.33 | 17.45 ± 1.28 | 14.49 ± 1.23 | 14.44 ± 1.12 | 10.89 ± 0.98 | 12.80 ± 1.03 | 6.78 ± 0.74 | 5.5 ± 0.54 | $F_{3,786}=0.70$, p=0.55 | $F_{3,790}=48.72$, p<0.0001 | $F_{1,789}=0.78$, p=0.38 |

Daily fecundity (DF) was significantly affected by host plants (*Χ*^2^_3_=129.33, p<0.0001), but not by *Wolbachia* (*Χ*^2^_1_=2.06, p=0.15) or its interaction with the plant (*Χ*^2^_3_=1.21, p=0.75; model 6; table 2). Contrast analyses revealed that DF was similar on purple and zucchini (3.37 ± 0.11 eggs per day; *contrast purple vs zucchini*: *Χ*^2^_1_=1.03, p=0.31), but higher on bean (4.60 ± 0.19 eggs per day; *contrast purple-zucchini vs bean*: *Χ*^2^_1_=40.14, p<0.0001), and lower on eggplant (2.10 ± 0.13; *Contrast eggplant vs purple-zucchini*: *Χ*^2^_1_=42.77, p<0.0001).

The effect of *Wolbachia* on egg hatching rate (HR) depended on the host plant tested (*Wolbachia*-plant interaction: F3,697=5.47, p=0.001; model 7; Table 1 and Fig. 2). Indeed, *Wolbachia* reduced HR on purple (F_1,172_=10.05, p=0.002) and on zucchini (F_1,177_=19.74, p<0.0001), but had no effect on bean and eggplant (F_1,181_=1.42, p=0.24 and F_1,158_=1.56, p=0.21, respectively).

**Figure 2.**
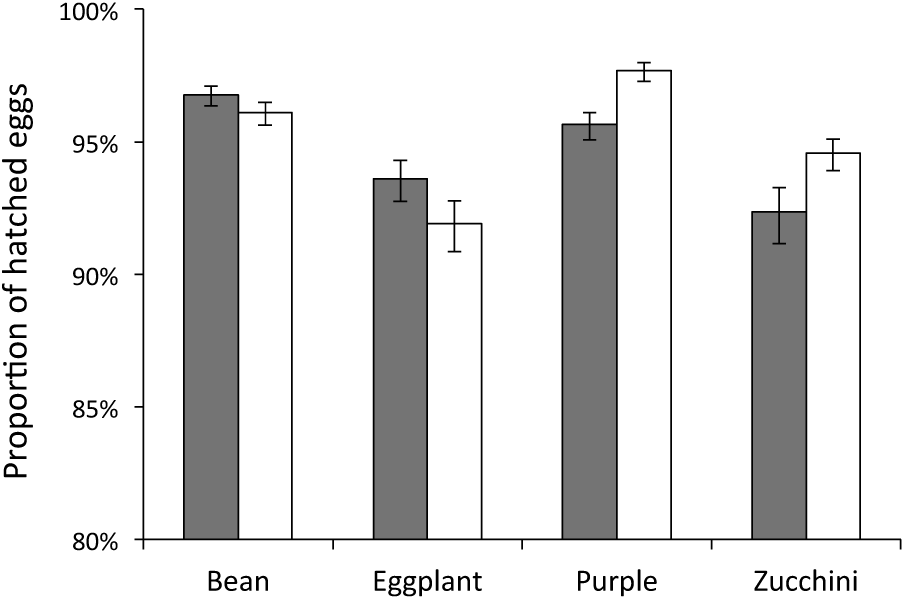
Effects of different host plants and of *Wolbachia* on the hatching rate of *T. urticae* eggs. Bars represent the mean (± s.e.) proportions of hatched eggs laid by *Wolbachia*-infected (Wi; grey bars) and uninfected (Wu; white bars) females on different host plants. Estimates were obtained from the GLMM statistical model that takes into account variation of fecundity among females, day within block as random effect, and corrects for zero-inflation. Standard errors were obtained from the upper and lower confidence intervals given by the model.

Juvenile mortality (JM) was not significantly affected by *Wolbachia* (F_1,692_=0.01, p=0.92; model 8; Table 2), and this was consistent across all host plants (*Wolbachia*-plant interaction: F_3,689_=1.85, p=0.14; model 8). However, host plant was a significant predictor of JM (F_3,693_=48.23, p<0.0001; model 8). Bean and zucchini did not differ significantly from each other (contrast bean vs zucchini: *Χ*^2^_1_=0.72, p=0.40) and led to intermediate JM of 16.8 ± 0.9%, while purple decreased it by 5.2 ± 1.5% (contrast purple vs bean-zucchini: *Χ*^2^_1_=53.82, p<0.0001), and eggplant increased it by 11.3 ± 2.1% (contrast bean-zucchini vs eggplant: *Χ*^2^_1_=109.36, p<0.0001).

*Wolbachia* infection affected differently the sex ratio (SR) produced on the different plants (*Wolbachia*-plant interaction: F_3,681_=2.48, p=0.04; model 9; Table 2 and Fig. 3). Indeed, *Wolbachia* decreased the proportion of males on purple (F_1, 168_=5.51, p=0.02) and on eggplant (F_1, 153_=8.54, p=0.004). On bean and zucchini, however, SR did not differ significantly between Wi and Wu mites (F_1,179_=5.51, p=0.54 and F_1,1726_=2.28, p=0.13, respectively).

**Figure 3.**
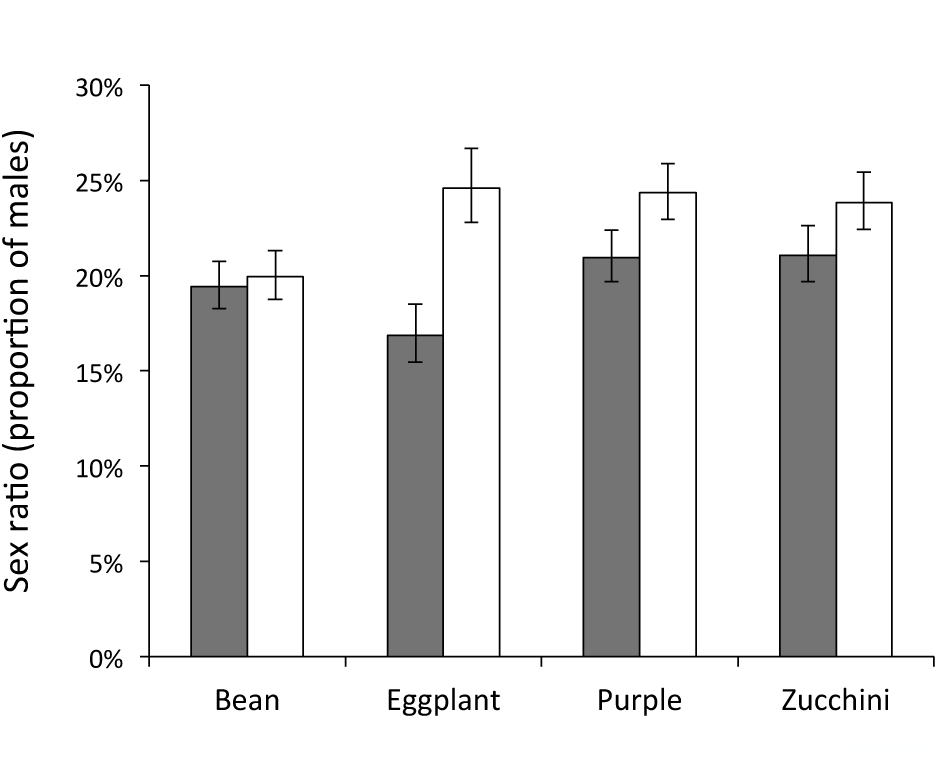
Effects of different host plants and of *Wolbachia* on the offspring sex ratio produced by *T. urticae* females. Bars represent the mean (± s.e.) proportions of male offspring produced by *Wolbachia-*infected (Wi; grey bars) and uninfected (Wu; white bars) females on different host plants. Estimates were obtained from the GLMM statistical model that takes into account variation of fecundity among females, day within block as random effect, and corrects for zero-inflation. Standard errors were obtained from the upper and lower confidence intervals given by the model.

Although we found a significant *Wolbachia*-plant interaction on HR and SR, *Wolbachia* did not significantly influence the average number of viable offspring (VO; F_1,789_=0.78, p=0.38), and this effect was independent of the host plant (*Wolbachia*-plant interaction: F_3,786_=0.70, p=0.55; model 10; Table 2 and Fig. 4). Nonetheless, host plant significantly explained this trait (F_3,790_=48.72, p<0.0001; model 10), with the highest values on bean, intermediate values on purple (*contrast purple* vs *bean*: *Χ*^2^_1_=4.82, p=0.03) and zucchini (*contrast zucchini vs purple*: *Χ*^2^_1_=5.12, p=0.02), and the lowest values on eggplant (*contrast eggplant zucchini*: *Χ*^2^_1_=44, p<0.0001).

**Figure 4.**
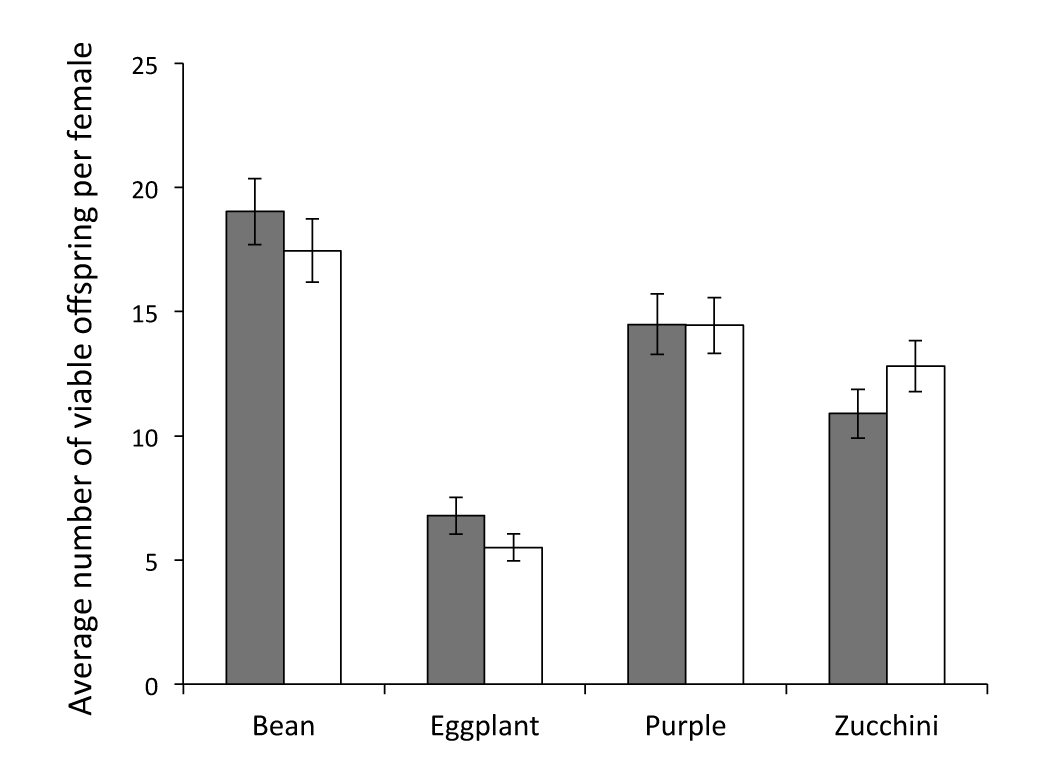
Effects of different host plants and of *Wolbachia* on the average number of viable offspring per female. Bars represent the mean (± s.e.) numbers of offspring (grey: sons; white: daughters) produced by *Wolbachia*-infected (Wi; grey bars) and uninfected (Wu; white bars) females on different host plants.

## DISCUSSION

In this study, we confirmed that *Wolbachia* is highly prevalent in *T. urticae* in Portugal, while *Cardinium* and *Rickettsia* were found at low prevalences (Zélé *et al.*, 2018). Moreover, this study suggests that endosymbiont prevalence varied with the host plant, *Cardinium* and *Rickettsia* being more prevalent on purple (although non-significantly for *Cardinium*) than on the other plants, and *Wolbachia* being more prevalent on bean and eggplant than on tomato, purple and zucchini. In the laboratory, *Wolbachia*-infected eggs had a lower hatching rate than uninfected ones on purple and zucchini, while this was not the case on bean and eggplant.

The prevalence of *Wolbachia* and *Rickettsia* in *T. urticae* females found in this study was relatively similar to that of an earlier study in the same geographical area (Zélé *et al.*, 2018). However, the prevalence of *Cardinium* was about five times lower in the current study than in the former one (2.5 ± 0.7 % *vs* 13.6 ± 2.9 %, respectively). As the populations were sampled on comparable host plants in this previous study (except for one population collected on *Datura stramonium*, the others were collected on bean, eggplant, tomato and zucchini), the discrepancy observed for the overall *Cardinium* prevalence between the two studies may be attributed to the time of collection. Indeed, mites were collected between September and December in the previous study and in June-July in the current one. Several studies have shown that the sampling period might affect endosymbiont prevalence and/or density in host populations (Toju & Fukatsu, 2011, Dorfmeier *et al.*, 2015, Martinez-Diaz *et al.*, 2016, Sumi *et al.*, 2017). This increase of *Cardinium* prevalence during summer is compatible with the hypothesis of an accumulation of this symbiont throughout the season via horizontal transfers (Zélé *et al.*, 2018).

We found that *Wolbachia* prevalence was overall high, but significantly higher on bean and eggplant than on the other plants. Whereas some earlier studies have shown that *Wolbachia* prevalence in herbivores varies according to the host plant (Ahmed *et al.*, 2010, Toju & Fukatsu, 2011, Guidolin & Consoli, 2017), including a recent study conducted in the spider mite T*etranychus truncatus* (Zhu *et al.*, 2018), others show no difference (Ji *et al.*, 2015). Unfortunately, the scarcity of studies, along with the fact that they were mostly done in other systems, hampers a meaningful comparison among studies. In addition, it is extremely difficult to sample spider-mite populations on all the plants tested within the same locality (see Table S1 in Additional file 1). Consequently, this implies an important sampling effort to obtain only a very reduced number of populations that fit the criteria for such studies. For instance, despite a large sampling effort across 21 localities and 12 host plant species, Zhu *et al.* (2018) could assess the effect of three common host plants (soybean, corn, and tomato) from three different locations only. Still, they did find that the prevalence of *Wolbachia* was significantly affected by the host plant (about 30% higher in tomato than in corn). In our study, the amplitude of the observed effects is much lower, possibly due to a threshold effect since the prevalence of *Wolbachia* that we observed in *T. urticae* is overall much higher than that observed in *T. truncates* by Zhu *et al.* (2018). Clearly, differences in *Wolbachia* prevalence were not associated with plant phylogenetic distance, as it differed between the solanaceous plants used (eggplant and tomato). Moreover, the effect of an endosymbiont on arthropod-plant interactions may depend on both the genotype (or species) of symbiont (Leonardo & Muiru, 2003) and arthropod host (Chen *et al.*, 2000, Ferrari *et al.*, 2007, McLean *et al.*, 2011, Wagner *et al.*, 2015), and/or their interaction (Ferrari *et al.*, 2007). More studies on plant-dependent symbiont prevalence may thus shed light on the potential factors underlying the pattern observed and on the ecological meaning of such effects.

Here, we hypothesize that the variation in endosymbiont prevalence according to the host plant is, at least partially, due to plant-specific effects of these symbionts on spider-mite performance. Although we did find some variation of *Rickettsia* and *Cardinium* prevalence according to the host plant, their prevalence was very low, so we opted for addressing this issue using *Wolbachia* only. Overall, we found a strong effect of the host plant on spider-mite performance, with the highest values observed on bean. This is not surprising, given that bean was the rearing environment of the population used, and is generally a host plant of high quality for spider mites (e.g. Magalhães *et al.*, 2011). Conversely, the lowest performances were found on Solanaceous plants (eggplant and tomato), being so low on tomato (cf. Material and Methods) that we excluded these data from further analyses. In the other four plants, we found that some traits (proportion escaping, female fecundity, and juvenile survival) were not affected by *Wolbachia* whereas others (egg hatching rate and sex ratio) were affected in a plant-specific manner.

The plant-specific effects of *Wolbachia*, although of low amplitude, could be explained by several non-exclusive mechanisms. First, *Wolbachia* may impose a nutritional burden to its hosts, sequestering and using vital host nutrients for its own survival (Chandler *et al.*, 2008, Caragata *et al.*, 2014, Ponton *et al.*, 2015), and this may vary with the host plant. Indeed, the nutrient composition of plant material is often poor or unbalanced for herbivores (Schoonhoven *et al.*, 2005, Karban & Baldwin, 2007), and nutrient deficient diet may increase the competition for resources between hosts and symbionts. In turn, this may lead to a decreased ability of infected spider mites to allocate enough nutrients to ensure egg viability on plants of low quality. Increased host-symbiont competition on such low-quality plants could also lead to a biased sex ratio towards males because females are produced from bigger eggs than males in *T. urticae* (Macke *et al.*, 2011). In addition, the slight *Wolbachia*-induced female-biased sex ratio observed on purple could be a consequence of the lower hatching rate observed on this plant, as larger eggs are generally more likely to hatch (Macke *et al.*, 2011). However, if this hypothesis would hold true, one would expect a stronger cost of *Wolbachia* in spider mites on plants of lower quality for mites, and we did not find such pattern.

Second, *Wolbachia* may directly influence the metabolism of some plants, which in turn can affect the biology of its herbivorous hosts. For instance, *Wolbachia* infecting the leaf-mining moth *Phyllonorycter blancardella* might be responsible for an increased level of cytokinins (plant hormones mainly involved in nutrient mobilisation and inhibition of senescence) in infested apple trees, *Malus domestica*. In this system, *Wolbachia* thus helps its host to develop in photosynthetically active green patches in otherwise senescent leaves (Kaiser *et al.*, 2010, Body *et al.*, 2013). Interestingly, cytokinins have also been shown to be responsible for sex-ratio shift towards females in the sap-feeding insect *Tupiocoris notatus* (although this effect was not mediated by *Wolbachia*; Adam *et al.* 2017). As *Wolbachia* possess a key gene involved in cytokinin biosynthesis in their genomes (Kaiser *et al.*, 2010), frequently infect the salivary glands of its hosts (Dobson *et al.*, 1999) and are present in high density in the gnathosoma of spider mites (Zhao *et al.*, 2013), one could speculate that the sex-ratio shift towards females observed in *Wolbachia*-infected mites on purple and eggplant in our study is mediated by increased cytokinin levels induced by *Wolbachia* in these two plants. Further research is thus needed to test this hypothesis. In particular, whether the *Wolbachia* present in spider mites also possess genes involved in cytokinin biosynthesis in their genomes is still unknown and the full genome of *Wolbachia* isolated from spider-mite hosts has, to our knowledge, not yet been sequenced.

Third, *Wolbachia* may interfere with the mites’ response toward plant defences. Indeed, endosymbionts found in herbivores, including *Wolbachia*, may directly manipulate the plant defenses to benefit their host (Frago *et al.*, 2012, Hansen & Moran, 2014, Zhu *et al.*, 2014, Sugio *et al.*, 2015, Giron *et al.*, 2017, Shikano *et al.*, 2017), or have a detrimental effect on their host by increasing the level of induced plant defences. For instance, down-regulation of several defense genes of maize by the western corn rootworm *Diabrotica virgifera* has been shown to be mediated by *Wolbachia* (Barr *et al.*, 2010, but see Robert *et al.*, 2013). Moreover, in a recent study, Staudacher *et al.* (2017) found that feeding by mites coinfected with *Spiroplasma* and *Wolbachia* increased the accumulation of 12-oxo-phytodienoic acid (a precursor of jasmonic acid) in tomato plants, compared to *Spiroplasma*-infected or non-infected mites. However, the concentration of jasmonic, salicylic and abscisic acids were not affected and no causal link could be established between the changes in plant defenses and mite performance (although only fecundity and longevity have been studied). Whether the presence of *Wolbachia* in *T. urticae* can upregulate the defences of zucchini and purple, and whether this could explain the reduced egg hatchability observed here, thus remains to be tested.

Despite the weak plant-specific effects of *Wolbachia* on mite performance, and that they do not affect the total number of viable offspring, they seem to be correlated with *Wolbachia* prevalence on field populations of *T. urticae* collected on different host plants. Indeed, given that *Wolbachia* is costly on egg hatchability on zucchini, we would expect a lower prevalence of this symbiont on this plant. Conversely, as *Wolbachia* increases the proportion of females produced on eggplant, we could expect a higher prevalence on this plant. Indeed, *Wolbachia* being maternally transmitted, it should always benefit from a more female-biased sex ratio. Note that, although *Wolbachia* may induce cytoplasmic incompatibility in *T. urticae* (Gotoh *et al.*, 2007, Xie *et al.*, 2011, Suh *et al.*, 2015), the effects observed in this study on spider-mite sex ratio cannot be attributed to this phenotype as it involves a cross between infected males and uninfected females, which was not performed here. On purple, we could expect the prevalence of *Wolbachia* to be intermediate, as the infection decreases egg hatchability but increases female proportion. Finally, bean being the plant on which spider mites have, overall, the best performance and that *Wolbachia* is not costly on this plant, we could expect its prevalence to be very high. Hence, by affecting the balance costs/benefits of *Wolbachia* on its spider-mite hosts, plants may affect *Wolbachia* prevalence. From the host perspective, however, although increased egg hatchability would probably benefit the spread of spider mites, it is not clear whether a female-biased sex ratio would benefit mites, as this is expected to depend on population structure (Hamilton, 1967, Macke *et al.*, 2011). More studies are thus needed to shed light on the potential role of *Wolbachia* on the host plant range of spider mites, as done in other systems (Hansen & Moran, 2014, Sugio *et al.*, 2015, Giron *et al.*, 2017).

In conclusion, our results show plant-dependent effects of *Wolbachia* on spider mites egg hatchability and offspring sex ratio, two crucial traits for both spider-mite population dynamics and *Wolbachia* spread among host populations. Although the amplitude of these effects is relatively low, they may, at least partially, explain the prevalence of this symbiont in spider mite populations collected on these different host plants. Moreover, our study highlights the importance of studying different host plants and life history traits when addressing the effects of endosymbionts on the performance of their herbivorous arthropods. These results also raised important questions, such as: (i) whether the pattern observed in this study varies between host and/or symbiont genotype, (ii) whether host plants affect the maintenance and/or spread of endosymbionts within and among populations, and (iii) whether endosymbionts affect the host range of herbivores.

## SUPPLEMENTARY DATA

Supplementary data are available at FEMSEC online.

## ACKNOWLEDGMENTS

We thank M. Bakırdöven, J. Denoyelle, L. Rodrigues, and Inês Santos for their help in spider-mite collection. We also thank IS for the maintenance of the plants and mite populations, Miguel Cruz, Nelson Martins, Jordi Moya Laraño and Susana Verala for advices in statistical analysis.

## AUTHOR’S CONTRIBUTIONS

Experimental conception and design: FZ, SM; field collections: JS, DG; acquisition of data: JS; statistical analyses: FZ, JS; paper writing: FZ, SM, with input from all authors. All authors have read and approved the final version of the manuscript.

